# The Brazilian Amazon Protected Area Network was largely unaffected by recent satellite-detectable fires

**DOI:** 10.1101/784975

**Authors:** Daniel P. Bebber

## Abstract

August 2019 saw dramatic increases in wildfires in the Brazilian Amazon, leading to arguments between Brazil and G7 leaders and widespread concern among conservationists. Popular media reports suggested that ‘swathes of the Amazon rainforest in Brazil are on fire’. Here we investigate the spatial distribution of fires through August 2019, showing that fires were largely restricted to deforested regions and areas with low canopy cover, particularly in unprotected areas that comprise half the region. In contrast, Brazil’s protected areas had one third as many fires, and forest in protected areas with high canopy cover was almost entirely unaffected by fire. Protected areas reduce deforestation and carbon emissions, and have proved largely untouched by recent fires. However, fires in closed-canopy forest cannot readily be detected by satellite remote-sensing and so this analysis likely underestimates the burned area in intact forest, both in protected and unprotected areas.

## Main text

Recent wildfires in the Amazon basin have caused widespread concern from conservationists, and have triggered arguments between the government of Brazil and G7 nations regarding responsibility for the protection of Brazil’s rainforests (1). The Brazilian Amazon is the world’s largest intact tropical forest, estimated to hold around 50 Gt of carbon in above-ground biomass (2). Deforestation rates in the Brazilian Amazon have varied widely over recent years with forest fires mainly affecting disturbed and logged areas (3), though droughts are becoming a greater trigger for fires (4). Fires are particularly severe during drought years which are often triggered by the El Nino Southern Oscillation (3–5). Low-intensity understorey fires under the canopy can lead to positive feedbacks, with more open canopy allowing dead trees to dry rapidly and fueling hotter and hotter burns (6). Preventing the establishment of such positive feedback loops is key to protecting remaining forest (3). Brazil has established numerous protected areas (PA) of different types and classifications, which form part of the world’s protected area network that covers around 15 % of the land surface (7). Deforestation rates are significantly lower within PAs than without (8), and tropical PAs are estimated to have reduced deforestation carbon emissions by one third in the first decade of the 21^st^ Century (7). Here we analyse the distribution of recent fires in the Amazon Basin to determine whether PAs have prevented forests from burning.

PAs cover approximately 45 per cent of the land surface of the Brazilian Amazon basin, with coverage scattered widely across the region (Fig. 1A). The southern border of the Brazilian Amazon is largely unprotected, however. Three quarters of the 645 PAs covering the basin are smaller than 4000 km^2^, while the largest at 96,500 km^2^ is the Yanomami Indigenous Area bordering Venezuela. PAs are highly forested, with 87 % of the area having a canopy coverage of over 90 % in 2018 at 0.01° spatial resolution (Fig. 1B). In contrast, only 46 % of unprotected areas have canopy cover over 90 % in 2018. Forest area loss was 38,814 km^2^ from 2000 to 2018 in PAs, compared with 306,394 km^2^ in unprotected areas, mostly along the ‘Arc of Deforestation’ along the south eastern border of the Amazon Basin (Fig. 1C). Forest area loss within PAs was greatest in the Triunfo do Xingu Environmental Protection Area in Pará (5652 km^2^ deforested), the Parque do Xingu Indigenous Area in Mato Grosso (2609 km^2^ deforested), and the Jamanxim National Forest in Pará (1551 km^2^ deforested).

**Fig. 1.**
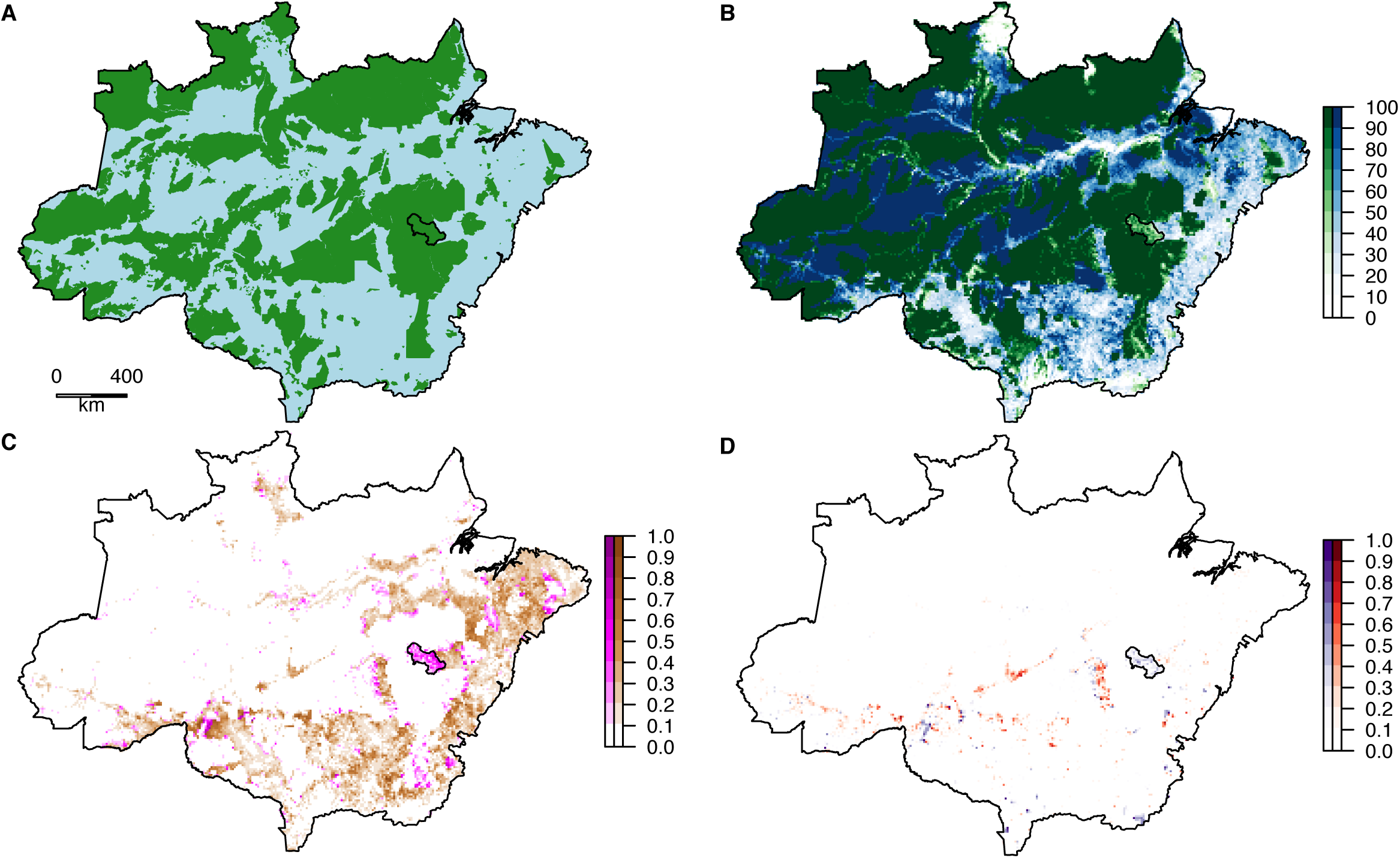
A. Brazilian Amazon with PA (green) and unprotected areas (blue). B. Forest canopy cover in 2018 (%) in PA (green) and unprotected areas (green). C. Deforestation fraction 2000-2018 in PA (magenta) and unprotected areas (brown). D. Fire incidence area fraction in August 2019 in PA (purple) and unprotected areas (red). The Triunfo do Xingu Environmental Protection Area is outlined in black (see text). Map data were aggregated from 0.01° to 0.1° resolution for plotting.

A record number of fires have been reported in the Brazilian Amazon in 2019, peaking in the middle of August(1). During August, fires were detected in a total of 18,005 km^2^ of PA forest and 58,954 km^2^ of unprotected forest, comprising 1.12 % and 3.19 % of the total PA and unprotected areas, respectively (Fig. 1D). The probability of fire detection varied with canopy cover in PA and unprotected areas (Fig. 2A). Assuming that areas deforested between 2000 and 2018 were effectively deforested (i.e. ignoring post-logging canopy recovery), in PAs the fraction of area with fires was greatest for mean canopy cover of between 20 and 30 %, while areas with more than 90 % canopy cover remained almost entirely unburned. In unprotected areas, the fraction of areas with fires was greatest for mean canopy cover between 60 and 70 %, again declining to near zero for forests with more than 90 % canopy cover. The fraction of area with fires increased with deforestation from 2000- 2018 in both PAs and unprotected areas (Fig. 2B). While overall the burned area was more than three times greater in unprotected areas than in PAs, the fraction of burned area increased more rapidly with deforestation in PAs than in unprotected areas.

**Fig. 2.**
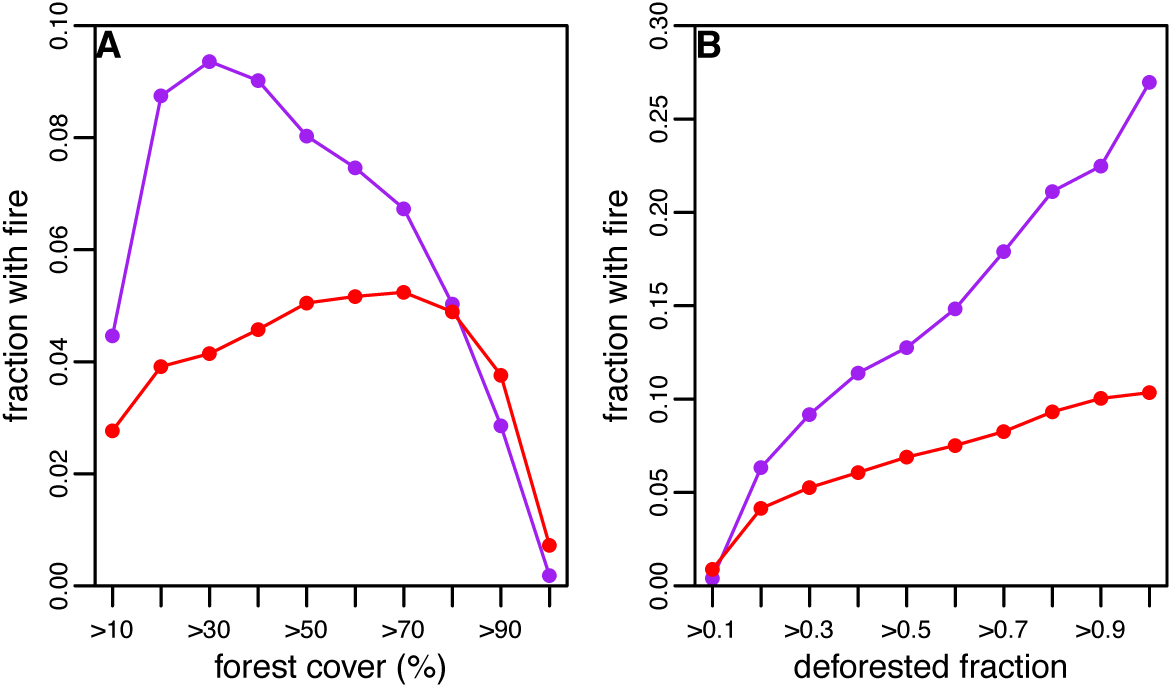
A. Fraction of area with fires by canopy cover class in PA (purple) and unprotected areas (red). B. Fraction of area with fires by deforestated fraction for PA (purple) and unprotected areas (red).

Fires were highly aggregated, occurring primarily in Pará (27,412 km^2^), Amazonas (15,074 km^2^), Mato Grosso (14,638 km^2^) and Rondônia (11,611 km^2^). Triunfo do Xingu Environmental Protection Area had the largest burned area (3676 km^2^ of a total area of 16,784 km^2^), comprising 20 % of all the burned area within PAs. Within Triunfo do Xingu, burned areas had lower canopy cover (interquartile range 18.5 – 73.8%) than unburned areas (interquartile range 40.2 – 98.0 %), and much greater levels of deforestation between 2000 and 2018 (interquartile range 0.23 – 0.81 vs. 0.00 – 0.54). Triunfo do Xingu was designated as a reserve in 2006, but between 2000 and 2018 mean forest cover fell from 96 % to 64 %. The reasons for this rapid rate of forest loss are unclear, but appear to be driven by illegal logging and cattle ranching (9). The small town of São Félix do Xingu (population approximate 45,000) lies on the south-eastern border of the reserve, across the Xingu River. Optical remote sensing images (Google maps) show large areas of agricultural land within the reserve, and so it appears that deforestation and forest fragmentation has permitted widespread fire. Other reserves with large burned areas include Jamanxim National Forest, Pará (1230 km^2^ burned), Jaci-Paraná, Roraima (1008 km^2^), Parabubure, Mato Grosso (915 km^2^) and Kayapó, Pará (739 km^2^).

There has been justifiable concern in the press and among politicians by the large areas of fire in the Brazilian Amazon during August 2019. Amazonian deforestation and fire emit around 0.2 Gt C per year (4), and threaten irreplaceable biodiversity (10). However, this analysis shows that the majority of satellite-detectable fires have occurred in unprotected areas, and that Brazil’s protected area network appears robust to fire outbreaks, particularly intact forest with high canopy cover and low rates of recent deforestation. Intact, undisturbed primary forests have lower fuel loads, higher relative humidity, lower maximum temperatures and lower vapour pressure deficit than secondary forests (11), and the clear conservation of canopy cover seen across most of Brazil’s protected area network is likely to have reduced fire frequency. Understorey fires under closed canopy, that are not detected directly by satellite sensors like MODIS and VIIRS (12), can lead to a positive feedback fire cycle with increasing burn intensity and canopy loss (6). The low rate of canopy loss in protected areas from 2000-2018 gives some hope that such positive feedbacks are not widespread within these undisturbed forests. While some reserves, like Triunfo do Xingu, have become highly degraded and vulnerable to burning in recent years, the recent fires show that protected area networks remain vital to reducing deforestation and fire in tropical forests.

## Methods

We obtained forest canopy estimates for the year 2000 and year of deforestation (2000 to 2018) at 0.0025 ° (approximately 30 m) resolution from the Global Forest Change database (13), available from http://earthenginepartners.appspot.com/science-2013-global-forest. We aggregated forest cover and deforestation data to 0.01 ° (approximately 1.2 km) resolution for computational efficiency and to match fire data resolution (see below). Forest cover in 2018 was estimated from the total deforestation fraction from 2000 to 2018, aggregated to 0.01°. The region of interest was the Brazilian Amazon Basin, defined by area of intersection of the Brazilian border (GIS polygon obtained from https://gadm.org) and the Amazon Basin (14) available from https://doi.org/10.3334/ORNLDAAC/1086. Forest and fire data were clipped to this region for analysis. We obtained protected area (PA) data from the WDPA (https://www.protectedplanet.net/), which has been used in previous analyses of the role of PAs in reducing deforestation carbon emissions (7). The GIS polygons for PAs were clipped to the Brazilian Amazon and only PAs in this region were considered in the analysis. The unprotected area was defined as the region within the Brazilian Amazon but not within a PA. We obtained fire observations from the MODIS and VIIRS systems for August 1^st^ to August 30^th^ 2019 from https://earthdata.nasa.gov/earth-observation-data/near-real-time/firms/active-fire-data. MODIS fire observations are given at 1 km resolution and VIIRS observations at 375 m resolution. The area burning within a pixel is not estimated, therefore fire observations were aggregated to 0.01° pixel resolution with any fire observation indicating a ‘burned’ pixel. Therefore, values for areas with fires are upper limits to the burned area, because actual fires may be smaller than the 0.01° pixel. All analyses were conducted in R version 3.6.1. GIS operations (e.g. aggregation of gridded data, intersection of polygons) were conducted using package *raster* version 3.0-2.

